# Water and phosphorus uptake by upland rice root systems unraveled under multiple scenarios: linking a 3D soil-root model and data

**DOI:** 10.1101/2020.01.27.921247

**Authors:** Trung Hieu Mai, Pieterjan De Bauw, Andrea Schnepf, Roel Merckx, Erik Smolders, Jan Vanderborght

**Author notes:** Joint first authors (Trung Hieu Mai, Pieterjan De Bauw).

## Abstract

**Background and aims:** Upland rice is often grown where water and phosphorus (P) are limited and these two factors interact on P bioavailability. To better understand this interaction, mechanistic models representing small-scale nutrient gradients and water dynamics in the rhizosphere of full-grown root systems are needed.

**Methods:** Rice was grown in large columns using a P-deficient soil at three different P supplies in the topsoil (deficient, suboptimal, non-limiting) in combination with two water regimes (field capacity versus drying periods). Root architectural parameters and P uptake were determined. Using a multiscale model of water and nutrient uptake, in-silico experiments were conducted by mimicking similar P and water treatments. First, 3D root systems were reconstructed by calibrating an architecure model with observed phenological root data, such as nodal root number, lateral types, interbranch distance, root diameters, and root biomass allocation along depth. Secondly, the multiscale model was informed with these 3D root architectures and the actual transpiration rates. Finally, water and P uptake were simulated.

**Key results:** The plant P uptake increased over threefold by increasing P and water supply, and drying periods reduced P uptake at high but not at low P supply. Root architecture was significantly affected by the treatments. Without calibration, simulation results adequately predicted P uptake, including the different effects of drying periods on P uptake at different P levels. However, P uptake was underestimated under P deficiency, a process likely related to an underestimated affinity of P uptake transporters in the roots. Both types of laterals (i.e. S- and L-type) are shown to be highly important for both water and P uptake, and the relative contribution of each type depend on both soil P availability and water dynamics. Key drivers in P uptake are growing root tips and the distribution of laterals.

**Conclusions:** This model-data integration demonstrates how multiple co-occurring single root phene responses to environmental stressors contribute to the development of a more efficient root system. Further model improvements such as the use of Michaelis constants from buffered systems and the inclusion of mycorrhizal infections and exudates are proposed.

## Introduction

Thirteen percent of world’s rice (*Oryza* spp.) cropping area is grown under upland conditions (GRiSP 2013; Chauhan *et al*. 2017). Upland rice systems are often used by subsistence farmers in Asia, Africa, and Central America and it serves as an important food source. Yield gaps of rice are largest in upland systems where drought events and phosphorus (P) deficiency form two major challenges for upland rice production, and these limiting factors often co-occur(Mueller *et al*. 2012; Diagne *et al*. 2013). For both water and P, root development and root characteristics are crucial for efficient resource acquisition under limiting conditions, and the performance of the rice roots is highly important to enable the plant cope with drought and/or P deficiency (Wissuwa and Ae 2001; Rose *et al*. 2013; Mori *et al*. 2016). Interestingly, soil water status and P availability are highly interrelated through diffusion, aeration, and sorption and both resources have specific behaviour and dynamics in the soil (Bünemann *et al*. 2011; Lal and Stewart 2016). Due to these complex interactions and the heterogeneous spatial distribution of both P (often stratified in top layers) and water (often more available in deeper layers) in soils, trade-offs and synergisms in P and water uptake efficiency of roots with respect to root architectural traits may exist (Ho *et al*. 2005). It is currently not known to what extent certain root types or root responses contribute to tolerance against droughts and/or low P availability and how root responses to limitations of one soil resource affect the uptake of another soil resource (i.e. water or P).

With experimental studies, we can evaluate the impact of P and/or water stress on the root development of rice. For example, an increased rooting depth and a decreased nodal root thickness were previously observed with reduced water availability, while a reduced root growth was observed in response to low P availability (De Bauw *et al*. 2019). Multiple studies have identified single root phene responses to environmental changes (e.g. Rose *et al*. 2013; Gao and Lynch 2016; Hazman and Brown 2018), but studies on root system responses integrating multiple phenes into a holistic root system performance are generally scarce. Single phene observations are however very important to identify interesting root parameters that influence P and water uptake, but an integrated understanding on the root system responses can only be assessed through mechanistic process modelling. The first step in such an approach is the integration of multiple root phene responses to environmental changes into one root system. To tackle this challenge, a dynamic structural-functional root model (i.e. CRootBox) was previously developed by Schnepf *et al*. (2018) enabling the integration of multiple root phenes. A second step would then be to link such integrated virtual root system with a soil-water model, in order to acquire insights in root performance and unravel the underlying processes of water and P uptake.

Underlying processes and mechanistic contributions of specific root responses to P or water uptake cannot be unravelled by experimental observations only and a more in-depth mechanistic understanding should be addressed by combining experimental results with coupled soil-root modelling (Lynch 2011; Ahmadi *et al*. 2014). Therefore functional-structural models on the interactions between root systems and soil are increasingly used to enhance physical understanding of nutrient transport and water flow in root-soil systems, along with experiments and analysis (Dunbabin *et al*. 2013). However, the major challenge in such approaches remained to capture rhizosphere gradients in the whole root system scale simulation and to transfer results from a single root scale to a larger scale efficiently and accurately. With this perspective, a continuum multiscale model that explicitly considers the 3-dimensional root architecture, water, and nutrient flow in both soil and roots as well as rhizosphere gradients around each root segment was recently developed by Mai *et al*. (2018).

Unfortunately, it remains very challenging to parameterize models so they can predict specific root responses of certain crops or specific varieties to environmental factors and models have to be informed with the reacting of particular plants to soil and environmental conditions. Combining single root phene analyses from real experiments with such 3D continuum multiscale soil-root models would theoretically enable to evaluate effects of water content, P sorption, and integrated root responses, and it would contribute to disentangling the underlying processes in the soil-root interface. Interestingly, such a combined approach was never thoroughly validated with real root data of upland rice, and we aim to highlight the potentials of combining single root phene analyses from real experiments with in-silico simulations by this 3D continuum multiscale soil-root model.

More specific objectives of this study were to (i) evaluate the integrated root system responses of upland rice to changing P and water availability; (ii) evaluate whether a 3D continuum multiscale soil-root model that is informed about the root development of rice can predict and explain water and P uptake from the soil under contrasting conditions; (iii) use such model to make predictions about the location of the water and P uptake; and (iv) evaluate the role of different root types and root characteristics for P and water uptake under contrasting scenarios in terms of P and water availability.

## Methods

### Laboratory experiments

#### Soil preparation, P treatments, and pot filling

A pot trial was set up (2017) in a greenhouse located at the Sokoine University of Agriculture in Morogoro (6°50’53.9”S, 37°39’31.3”E; Tanzania). The averages of the daily minimum and maximum temperatures were respectively 21.9 °C and 33.4 °C. Initially, a P-deficient soil (0.035 mg P l^−1^ in soil solution, measured by ICP-MS after a water extraction) was collected from an upland rice field in Matombo (07°02’46.8”S; 37°47’11.6”E; Tanzania). This soil was classified as a ferralsol (*World Reference Base for Soil Resources*) and was characterized by a soil pH(CaCl2) = 5.7, a particle size distribution of 9% sand, 57% silt, 34% clay, and an oxalate extractable Alox = 1073 mg kg^−1^, Feox = 1730 mg kg^−1^, Mnox = 2559 mg kg^−1^, Pox = 122 mg kg^−1^. After sampling, the bulk soil was shade dried, crushed to an aggregate size of 4 mm, and amended with salts of NH4NO3, KCl, CaCl2, MgSO4, ZnSO4, CuSO4, H3BO3 and Na2MoO4 at rates of 37 mg N kg^−1^, 95 mg K kg^−1^, 16 mg Mg kg^−1^, 21 mg S kg^−1^, 3.5 mg Zn kg^−1^, 0.04 mg B kg^−1^, 0.08 mg Cu kg^−1^, and 0.03 mg Mo kg^−1^ soil, in order to avoid any deficiency other than P.

As P generally accumulates in the topsoil, no P was initially added to the bulk soil in order to mimic a P deficient subsoil. Large pots (height: 55cm, diameter: 16cm) were then filled with 7.3 kg of the P deficient subsoil. The remaining of this bulk soil was then subjected to three different P treatments. One third was amended with a non-limiting amount of ground Triple Super Phosphate (TSP) (70.8 mg P 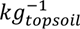 or 354 mg P per pot) up to a theoretical P concentration of 0.5 mg P l^−1^ in soil solution (PlusP), which was based on the previously determined P adsorption isoterm. Another third was amended with a sub-optimal amount of ground TSP (25.0 mg P 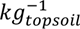 or 125.2 mg P per pot) up to a theoretic P concentration of 0.1 mg P l^−1^ in soil solution (SubP). The remaining part was not amended with TSP (NoP), and CaCl2 was used to equalize the amended Ca in all treatments. Pots were then filled with 5 kg of topsoil, affixed on top of each subsoil. The bulk density of the dry soil in the pot was 1.29 g cm^−^³. The layer of the subsoil was 30 cm and the topsoil was 20 cm thick. Pots were then irrigated to bring the whole pot to field capacity (38% w/w).

#### Sowing, maintenance, and water treatments

One pre-germinated seed of the typical upland rice variety (NERICA4) was sown into the pots (1 cm depth) at the center of the surface, which closely relates to a conventional spacing density of 20×20 cm for rice. NERICA4 is developed by the Africa Rice Center using interspecific crosses between *Oryza sativa* (Asian rice) and *Oryza glaberrima* (African rice). NERICA4 is an upland rice variety known for its drought tolerance but it is relatively susceptible to low P.

Two top dressings of NH4NO3 (in solution) were later applied at a rate of 349 mg N per pot at 21 and 34 days after sowing (DAS). An additional top dressing of ZnSO4, ZnCl2, KCl, and MgSO4 were added to each pot at rates of 0.27 g Zn, 0.58 g K, 0.16 g S, and 0.11 g Mg.

Pots were daily irrigated to field capacity (based on pot weight) until 25 DAS, and two contrasting water treatments were then initiated and maintained until the end of the trial. Half of the pots were daily irrigated to field capacity (FC), while the other half was subjected to drying periods (DP). In order to represent drying cycles during erratic rainfall, pots were re-watered up to field capacity after a period of ca. six days (preset as an average period of drying). Each treatment combination was replicated four times. The amount of irrigated water was consistently monitored to assess evapotranspiration, water use, and water productivity. An estimate of the evaporation was monitored by daily weighing and re-irrigating six unsown pots, randomly placed through the experiment.

#### Data collection

Plant development was monitored by measuring plant height and counting tillers and leaves twice a week. At 52 DAS, shoots were cut, oven dried (60 °C), weighed, and manually ground by mortar. P concentrations in the shoot tissues were then determined by ICP-OES (Thermo Scientific iCAP 7000 series) after digestion in HNO3.

Immediately after removing the shoot, the soil cylinder was carefully taken out of the pot and precisely cut into three segments. One part comprised a segment (A) from 0 to 15 cm depth which included the ‘*shallow roots*’; another segment (B) comprised soil from 15 to 30 cm depth including the ‘*intermediate roots*’; and the last segment incorporated the ‘*deep roots*’ below a depth of 30 cm. The latter segment (C) was defined according to most rice studies, where deep roots are defined as roots below 30 cm (Kato *et al*. 2006, 2013; Gowda *et al*. 2011). For each soil segment, roots were carefully washed out by gently shaking the soil segments on a 2 mm net in water. After removing the soil from the roots, roots of each segment were placed in a dish with clean water and root architectural variables were determined as follows.

The number of nodal roots was counted and the average nodal root diameter was measured using a transparent ruler (0.1 mm). For each segment the transparant ruler was placed on the nodal roots, and the average thickness was visually determined. S-type lateral root density (i.e. the branching distance of the first order laterals on the nodal root) was scored using the ‘shovelomics scoreboard’ developed by Trachsel et al. (2011). The shovelomics scoreboard is a resource for phenotyping roots of soil-grown crops after excavation. S-type laterals are short and thin lateral roots, emerging at the root base on the nodal roots and they don’t have higher order branches. L-type laterals are longer and generally thicker, and they branch further into higher order branches (Nestler *et al*. 2016). The density of the S-type laterals was determined by placing the different scoring classes from the scoreboard next to the roots and comparing the densities from the board with the actual density on the root. These scores were then translated into actual values of distance. Lateral root thickness (both at the base and at the deeper roots) were visually scored according to five classes, each corresponding to a thickness class with actual diameter values. The secondary branching distance on L-type roots was scored also based on the ‘shovelomics scoreboard’. After analyzing root architecture in the different segments, the roots from each segment were oven dried (60°C) and weighed to determine root distribution and biomass allocation. Total P uptake was calculated by assuming an equal P concentration in root and shoot.

### The continuum multiscale model for growing root systems and virtual experiment setup

#### General model description

The CRootBox root architectural model is a tool to create root geometries based on root growth mechanisms such as elongation, branching, and death. The root is represented as a 1-dimensional branched network discretized into connected line segments with attributes such as root radius and age. Those attributes may be accessed in simulations using this root grid and may for instance affect root hydraulic properties. In this work, the root architecture model parameters were obtained by calibrating the model based on the available data from the lab experiment.

The resulting root grid was used in a numerical simulator, DuMu^x^, in which the 1-dimensional root network is embedded within a 3-dimensional soil domain. Water flow and solute transport are solved in both domains, and exchange of water and solutes between domains is modelled based on source/sink terms. Potential transpiration and irrigation events are the main drivers for water flow, they are prescribed as boundary conditions at the soil surface and root collar, respectively. The main factors influencing P uptake are the heterogeneous distribution of P concentration in the soil together with the root uptake parameters as prescribed by the Michaelis Menten kinetics for each root segment. In addition, our multiscale approach handles small-scale concentration gradients by assigning a 1-dimensional radially symmetric rhizosphere model to each root segment (Mai *et al,* 2018). It combines the Barber-Cushman model approach for single roots with the 3-dimensional macroscopic model approach in a mass conservative way.

Root growth is mimicked by considering, at each time step, only those root segments that have already been borne. If a root enters a soil control element that already contains other root segments, the zone of influence of each root segment is re-computed in a way that assigns a new soil cylinder with homogeneous concentration to the new root segment while the domains of the already existing root segments are clipped from the outer end, keeping the already existing rhizosphere concentration profiles. The overall aim is to compute overall water and P uptake in the correct order of magnitude for different scenarios (comparable with measurements) and to use this model to investigate the dynamics and aspects on P & water uptake that can not be measured in the real experiment so as to obtain a deeper understanding the system, e.g. to determine the key drivers, locations, and root types important for water and P uptake.

#### The mathematical details and their implementation

To simulate the water and nutrient transport in the soil-rice columns, the multiscale model developed by Mai *et al*. (2018) was employed and further developed to consider root growth and a dynamic root system. The model considers two domains: the soil and the root system. The root system is described by a set of connected linear root segments (their length is in the order of mm to cm and defines the spatial resolution along the root axis). In the soil domain, two spatial scales are considered: the macro and rhizosphere scale. The macro-scale model simulates soil water flow and nutrient transport at a spatial resolution in the order of one cm. The rhizosphere scale captures the steep radial nutrient gradients from the root surfaces in 1D cylindrical models around single root segments and is used for calculating the resource uptake in sub-mm resolution. The water flow in the soil domain is described by the Richards equation (Richards 1931) and the water flow in the 3D root system is solved using the approach developed by Doussan *et al*. (2006). The flow through these two domains are coupled via source/sink terms which depend on the water potentials in the root xylem and at the root-soil interface. The nutrient transfer in the soil is described by the transport equation with consideration of nutrient sorption onto the soil matrix, and diffusive and advective transport. The advective transport is directly coupled to the soil-plant water flow model whereas the diffusive transport is indirectly coupled through the water content dynamics which influence the effective soil diffusion coefficient. The sorption-desorption process is assumed to be in a dynamic equilibrium which is described by a non-linear (Freundlich) isotherm that was derived from batch sorption experiments. So far, we did not consider kinetic sorption-desorption or multi-species sorption-desorption processes or other bio-geochemical reactions in the rhizosphere which can be influenced by root exudation. Nutrient uptake from the soil per unit root surface area was described as a function of the nutrient concentration at the soil root interface using Michaelis Menten kinetics. The Michaelis Menten parameters were assumed to be constant and the same for all root segments. The root development was simulated using the root architecture model CRootBox (Schnepf et al., 2018). The root system of rice was characterized as having several nodal roots and two types of lateral roots: the short and thin laterals (S-type) versus the longer ones (L-type) as discussed in the previous section. CRootbox was used to generate six virtual plants, one for each P-water treatment, with similar features as the observed root systems (e.g. lateral density, lateral thickness, number of nodal roots, nodal thickness, root mass distribution along depth,…). More details about the water flow and nutrient transport models, the implementation of the dynamic root growth in the flow and transport model, the root growth model, the mathematical equations, and the multiscale coupling method are presented in Supplementary Information (Text S1) and can be found in Mai *et al*. (2018). The model code is shared on GitHub (https://github.com/Plant-Root-Soil-Interactions-Modelling/dumux-rosi/tree/pub/Mai2019).

#### Virtual experiment setup

Based on the data from the lab experiment, simulation scenarios were set up to study the water and P transport in the soil-rice column as well as the functions of the root system in plant water and nutrient uptake. Two water regimes were considered in the model: (i) optimal conditions where the soil water content was maintained at field capacity (FC) by daily compensating evapotranspiration, and (ii) conditions with drying periods (DP) where the soil columns were not irrigated for several days. Similar to the lab experiment, P availability in the simulations comprised three contrasting concentrations in the topsoil: P deficient (NoP), a suboptimal (SubP), and a non-P limited scenario (PlusP). Combining these three P rates and two water regimes resulted in six contrasting scenarios to be simulated.

##### Reconstruction of the soil column

The numerical mesh to represent the soil column domain, which has a diameter of 16cm and 55cm length, was generated using the mesh generator GMesh (Geuzaine and Remacle 2009). The soil column was divided vertically into 2cm layers. Each soil layer was discretized to tetrahedral elements with 16 cubic elements at the centre. In total, the soil column has 1839 elements. The soil column is considered as a homogenous medium with constant hydraulic properties and sorption capacity for P which is characterized by the Freundlich isotherm. The Freundlich coefficient and the Freundlich power were derived from a P adsorption experiment. Briefly, the soil was dried and sieved. Replicate samples of 3 g were suspended in 30 mL water and amended with KH2PO4 at various rates between 0us ratel^−1^. The soils were equilibrated with an end-overend shaker for 24 h followed by centrifugation and filtration (0.45 μm) and analyzed of P by Inductively Coupled Plasma Mass Spectrometry (ICP-MS, Agilent7700X). The Freundlich coefficient was 124.8 mg^0.6^ kg^−1^ L^0.4^ and the Freundlich Power was 0.4. The soil parameters values are presented in Table 1, and these were derived from direct measurements or from parameter values that were available in literature. Regarding the distribution of the P concentration in the soil column, the P concentration in the topsoil layer (the first 20 cm layer from the soil surface) varied according to the P rate, while the P concentration in the deeper subsoil (from 20cm to the bottom) remained constant and deficient.

**Table 1:**
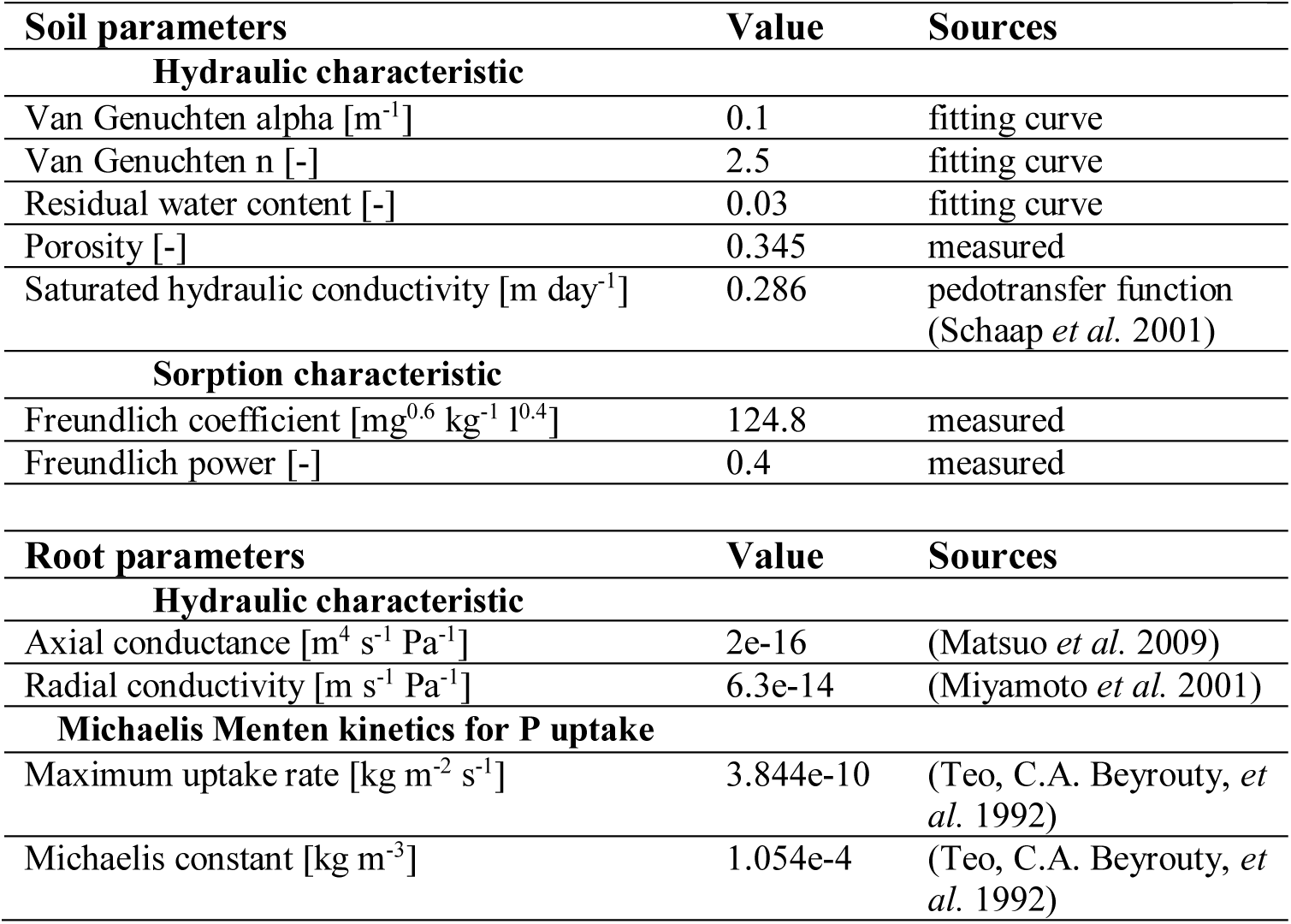
Soil and root parameters in the multiscale simulations

##### Reconstruction of rice root systems

From the experimental data, we generated the rice root systems by using CRootBox. The rice root systems consist of several nodal roots and a large number of lateral roots. The number of nodal roots varied from 43 to 100 depending on the water and P conditions during the growth. The root parameters used in CRootBox are presented in Table 2. To parameterize the root system architecture in each scenario, the average of each measurement for the different plant replicates (i.e., lateral root density, S-type and L-type lateral root radius, and secondary branching distance on L-type roots) were used as input parameters (Table 3). The radius of the second order lateral on L-type laterals was assumed to be the same as the radius of the S-type roots. We used the measured root mass fraction in each soil segment and the total root mass to calibrate the root model as illustrated by Figure S1 (supplementary information). The emergence coefficient *k* of the nodal roots, the transition depth *z*_0_, and the scaling factor of the interbranch distance at the bottom of the soil column *Ks* were adjusted to fit with the root mass distribution derived from the experiment (Text S1, Supplementay Information). The number of root segments in the root system varied between 13633 segments in case of stresses of water and P and 27659 segments in case of optimal water condition (FC) and surplus P in top soil. For the simulation of water and nutrient uptake, the hydraulic root parameters and Michaelis Menten parameters are taken from literature and presented in Table 1

**Table 2:** CRootBox parameter values used for the reconstruction of the rice root system of NERICA4 grown under contrasting conditions. P availability comprised three levels (NoP, SubP, PlusP) while water included two levels (field capacity (FC) versus drying cycles (DP)).

|  | -----FC----- |  |  | -----DP----- |  |  |
| --- | --- | --- | --- | --- | --- | --- |
|  | NoP | SubP | PlusP | NoP | SubP | PlusP |
| **Nodal roots** |  |  |  |  |  |  |
| Emerging coefficient* [day<sup>-1</sup>] | 12 | 8.5 | 13.5 | 10 | 5 | 12.5 |
| Maximal root length [cm] | 90 | 90 | 90 | 90 | 90 | 90 |
| Initial elongation rate [cm day<sup>-1</sup>] | 2 | 2 | 2 | 2 | 2 | 2 |
| Length of basal zone [cm] | 1 | 1 | 1 | 1 | 1 | 1 |
| Length of apical zone [cm] | 1 | 1 | 1 | 1 | 1 | 1 |
| Insertion angle [rad] | 0.7 | 0.8 | 0.8 | 0.7 | 0.8 | 0.8 |
| Tropism type [-] | 1 | 1 | 1 | 1 | 1 | 1 |
| Tropism strength [-] | 0.7 | 0.7 | 0.7 | 0.7 | 0.7 | 0.7 |
| Root flexibility [cm<sup>-1</sup>] | 0.2 | 0.2 | 0.2 | 0.2 | 0.2 | 0.2 |
| **S-type lateral roots** |  |  |  |  |  |  |
| Maximal root length [cm] | 1.5 | 1.5 | 1.5 | 1.5 | 1.5 | 1.5 |
| Initial elongation rate [cm day<sup>-1</sup>] | 1 | 1 | 1 | 1 | 1 | 1 |
| Tropism type [-] | 1 | 1 | 1 | 1 | 1 | 1 |
| Tropism strength [-] | 3.5 | 3.5 | 3.5 | 3.5 | 3.5 | 3.5 |
| Root flexibility [cm<sup>-1</sup>] | 1 | 1 | 1 | 1 | 1 | 1 |
| **L-type lateral roots** |  |  |  |  |  |  |
| Maximal root length [cm] | 6.5 | 4 | 4 | 6.5 | 4 | 4 |
| Initial elongation rate [cm day<sup>-1</sup>] | 1.5 | 1 | 1 | 1.5 | 1 | 1.5 |
| Length of basal zone [cm] | 0.5 | 0.5 | 0.5 | 0.5 | 0.4 | 0.37 |
| Length of apical zone [cm] | 0.5 | 0.5 | 0.5 | 0.5 | 0.4 | 0.37 |
| Inter-branch distance [cm] | 0.7 | 0.7 | 0.6 | 0.5 | 0.4 | 0.37 |
| Insertion angle [rad] | 1.2 | 1.2 | 1.2 | 1.2 | 1.2 | 1.2 |
| Tropism type [-] | 1 | 1 | 1 | 1 | 1 | 1 |
| Tropism strength [-] | 3.5 | 3.5 | 3.5 | 3.5 | 3.5 | 3.5 |
| Root flexibility [cm<sup>-1</sup>] | 0.2 | 0.2 | 0.2 | 0.2 | 0.2 | 0.2 |
| **Second order lateral roots** |  |  |  |  |  |  |
| Maximal root length [cm] | 0.8 | 0.8 | 0.8 | 0.8 | 0.8 | 0.8 |
| Initial elongation rate [cm day<sup>-1</sup>] | 1.5 | 1 | 1 | 1.5 | 1.5 | 1.5 |
| Tropism type [-] | 1 | 1 | 1 | 1 | 1 | 1 |
| Tropism strength [-] | 1 | 1 | 1 | 1 | 1 | 1 |
| Root flexibility [cm<sup>-1</sup>] | 0.2 | 0.2 | 0.2 | 0.2 | 0.2 | 0.2 |
| **Lateral roots distribution** |  |  |  |  |  |  |
| Lateral Type Transition Steepness* [cm<sup>-1</sup>] | 0.3 | 1.5 | 1.5 | 0.3 | 0.3 | 1.5 |
| Lateral Type Transition Depth* [cm] | -12 | -20 | -20 | -15 | -18 | -15 |
| Inter-Branch Distance Scaling Factor* [-] | 1.6 | 1.85 | 2.3 | 1.8 | 1.9 | 1.65 |
| Inter-Branch Distance Slope* [cm<sup>-1</sup>] | 1.5 | 1.5 | 1.5 | 1.5 | 1.5 | 1.5 |
| Inter-Branch Distance Transition Depth* [cm] | -13 | -20 | -20 | -25 | -15 | -20 |

**Table 3:**
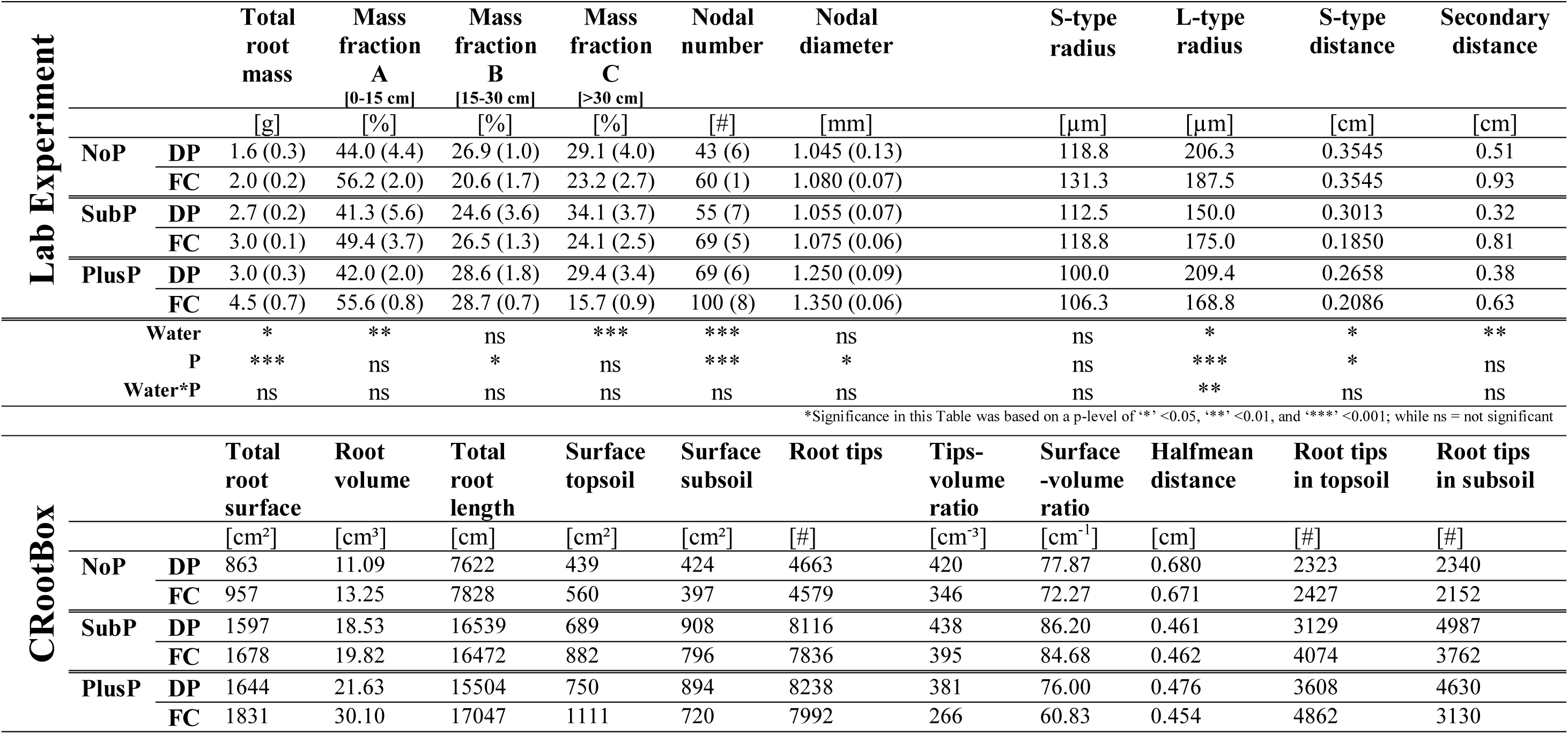
Rice root parameters determined in the lab experiment (top) and derived from CRootBox (bottom). Root architectural parameters extracted from the lab experiment (top) were used as an input to develop the virtually growing 3D root systems in CRootBox. The mass fraction distribution measured in the lab experiment was used to simulate equal volume fractions in the corresponding layers. Rice was grown on a P deficient soil with three P treatments in the topsoil (No P amendment (NoP), a suboptimal rate (SubP), and a non-limiting rate (PlusP)) and two water levels (Field Capacity (FC) and Drying Periods (DP)). S- and L-type radius and the distance among S-types and secondary branches have been estimated based on the average of the root scores (Table S1, Supplementary Information).

##### Top boundary condition and initial P concentrations

In order to simulate the water conditions in the virtual experiment, the daily irrigation data as well as the evaporation rate monitored in the real experiment was used as the boundary condition in the soil surface (Figure S2). The daily evaporation rate varied in the range of 18-120 mL day^−1^. In order to keep the soil column saturated in the FC treatment, the irrigation rate compensated the evaporation and transpiration of rice plant which increases over time following plant development. For the DP scenario, four cycles of drying (no irrigation) were reproduced from the 25^th^ day of the experiment by virtually withholding irrigation. Total transpiration in each scenario was derived by extractiong the monitored irrigation by the measured evaporation. Daily transpiration rate was then calibrated proportional to the growing number of leaves. Evapotranspiration rates per unit surface calculated in this trial were relatively high, which probably attributes to the fact that the rice canopy exceeds the pot surface. At the bottom and sides of the soil column, zero-flux boundary conditions were imposed. In the subsoil, the initial P concentration was 0.035 mg P l^−1^ for all scenarios. In contrast, the initial P concentration in the topsoil was varied corresponding to the P availability in solution after the three P fertilization levels in the real experiment: 0.035 mg P l^−1^ (NoP), 0.1 mg P l^−1^ (SubP) and 0.5 mg P l^−1^ (PlusP).

## Results

### Evaluating the rice root responses to changing P and water availability, and insights revealed by CRootBox

The shoot and root biomass significantly responded to the P and water treatments with a significant interaction term (details not shown). The same was true for total (root+shoot) P uptake that significantly increased with increased P supply and increased water supply (Figure 2) and there was a significant treatment interaction on P uptake. That interaction shows larger effect of water regime on P uptake at higher P supply and no effect of water regime under P deficient conditions (Figure 2).

Table 3 presents the observed root responses of upland rice to combinations of contrasting P rates and subjected to field capacity (FC) and drying periods (DP). Data from the pot experiment reveal that the total root mass increased with increasing P rates and decreased under DP compared to FC. Both the absolute root mass and the mass fraction in the shallow layer (0-15 cm) decreased with DP (Table 3 & Figure S1, supplementary information). The mass fraction in the intermediate layer remained unaffected by water but increased with increasing P rate. The absolute root mass in the deep layer (>30cm) remained unaffected by the water regime, but the root mass fraction (% of total root mass) in this deep layer systematically increased under DP. (Table 3 & Figure S1)

Data from the pot trial show that the number of nodal roots was largest under PlusP and it decreased with DP. Nodal thickness was largest under PlusP. It showed a decreasing trend under DP, but this was not significant. The density of S-type laterals was smallest under NoP, and it generally decreased under DP, while the density of the secondary roots strongly increased under DP. The L-type radius under drought was smallest for SubP compared to NoP and PlusP. (Table 3)

Root mass distribution in the soil column was calibrated in CRootBox with a good agreement based on the observed distribution in the pot experiment (Figure S1). Following these distributions, the interbranch scaling factor and the calibrated transition depth of the L-type laterals were derived (Figure S3 & S4; supplementary information). The 3D structure of the contrasting root systems and the distribution of the root types (simulated in CRootBox) grown in contrasting environments are visually presented on Figure S5 (supplementary information), while Figure 1 illustrates the reconstructed 3D architecture of these root systems with the corresponding P uptake rates by the root system. The 3D root systems reveal that the total root surface and root volume generally increased with increasing P rate and water availability (bottom of Table 3). A similar increase with P rate is true for root length under FC, but not under DP. 3D simulations reveal that under DP, the root surface in the topsoil generally decreased compared to FC, while the root surface (and root length) in the subsoil increased. The total number of root tips (i.e. the terminal portion of each root type including S-type laterals, L-type laterals, secondary laterals, and nodal roots) generally increased with increasing P rate, and it also increased under DP compared to FC, which mainly follows an increase in root tips on the L-type roots under DP (data not shown). Under DP, the number of root tips in the subsoil exceeded the number of root tips in the topsoil, while the reverse was true under FC. (Table 3)

**Figure 1:**
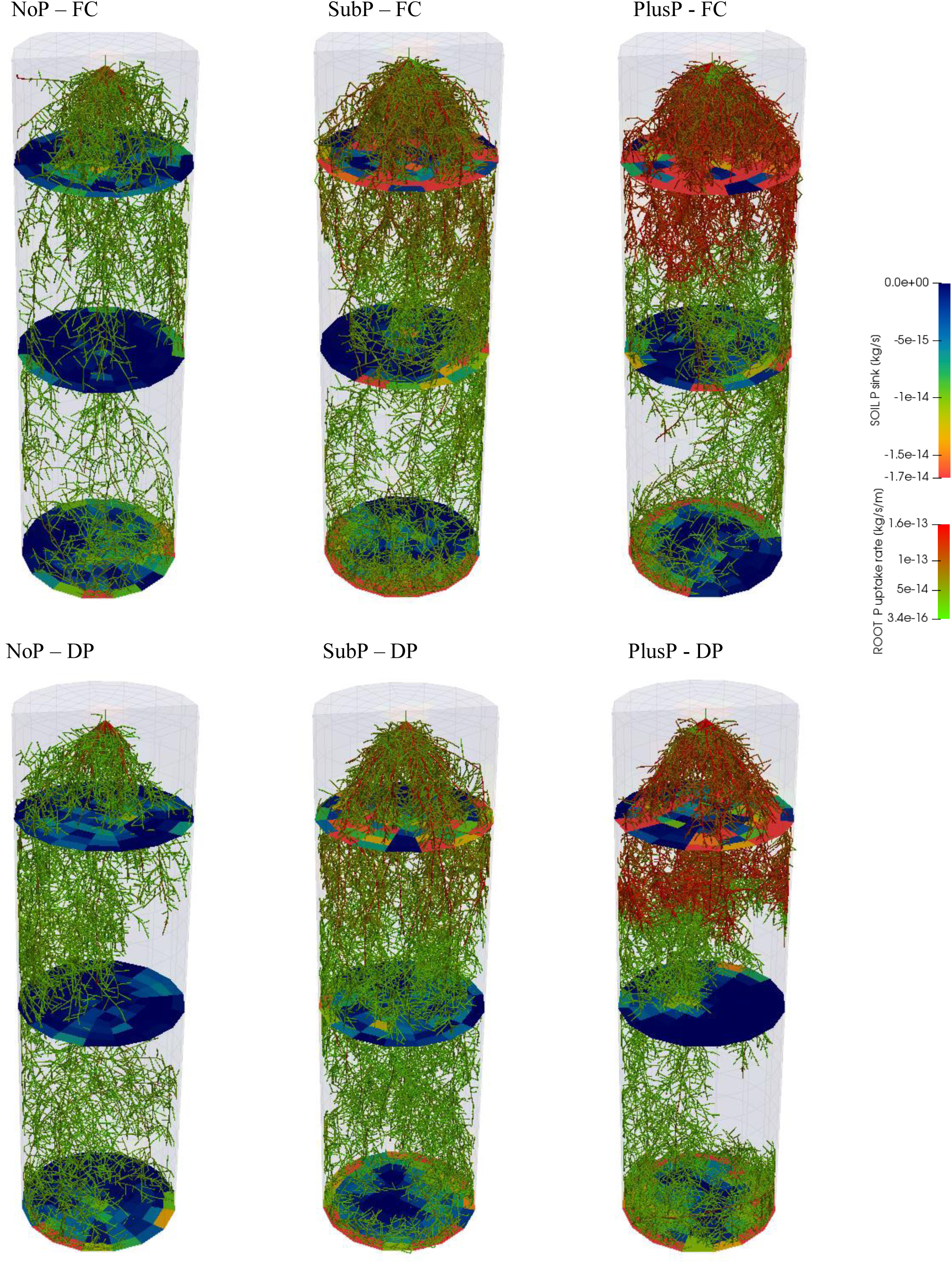
Simulated root systems (3D architture) from CRootBox of upland rice grown on a P deficient soil with three P treatments in the topsoil (No P amendment (NoP), a suboptimal rate (SubP), and a non-limiting rate (PlusP)) and two water rgimes (Field Capacity (FC) and Drying Periods (DP)). The color of the root presents the P uptake rate of the root, while the color on the discs present the soil P sink. The transition between topsoil and subsoil (at 20 cm depth) can be observed by the shift in P uptake rate in SubP and PlusP.

The surface-volume ratio (i.e. the ratio of the root surface to the volume of the root system) and root tip-volume ratio (i.e. the ratio of the total number of root tips to the total volume of the root system) were generally largest under SubP, and it increased under DP compared to FC within each P treatment (Table 3). The surface-volume ratio generally reached its maximum before 10 DAS and it subsequently decreased with growth. Simulations further reveal that this root surface per root volume was consistently largest for the L-type laterals, followed by S-type laterals, and being smallest for nodal roots (data not shown). The half mean distance was largest under NoP due to its low root density in the soil column and this value decreases under PlusP (Table 3).

Analysis of the reconstructed root systems further shows that under DP, the total contribution of L-type roots to both root mass and root surface increased compared to FC, while the contribution of S-type laterals decreased. The total root surface comprised by the L-type laterals (which include the secondary laterals on these L-types) was generally larger than the surface comprised by S-type roots, but S-type roots are more located in the upper layer. Interestingly, the root surface of L-type roots in the topsoil dramatically increased under DP, while the root surface of the S-type roots in the topsoil decreased. (data not shown)

### Simulated water dynamics and effects on P diffusion

The change of the water content in the soil column during the drying cycles in the DP treatments is shown on Figure S10 (supplementary information). Under FC, the water content kept relatively constant at saturated state (despite some minor daily fluctuations). During thedrying cycles, plants continue to transpire although irrigation was stopped. therefore the water content dropped about 10 to 20%. The soil water content dropped more strongly when higher P levels were applied in the topsoil. It’s also observed that the effective P diffusion at root surface under FC generally remained constant with minor daily fluctuations (Figure S11, supplementary information). In contrast, under DP, the effective P diffusion coefficient dramatically decreased 30-50% during the drying cycles.

### Evaluating simulated versus measured P uptake

Figure 2 presents the simulated P uptake by the reconstructed 3D root systems through the multiscale model versus the averages of the measured P uptake in the lab experiment. Both the simulations and the actual P measurements show that the total P uptake strongly increased with increasing P rate. Under DP both the measured and simulated P uptake is reduced in the PlusP and SubP scenario, but this was not observed for NoP. The negative effect of DP on total P uptake thus increased with increasing P rate in both simulations and actual measurements. At the first glance, the P uptake from the 3D virtual experiments is close to the measured values from the real experiment. However, the simulated P uptake was underestimated for NoP (ca. 50% underestimation), while the simulations slightly overestimated the P uptake for PlusP under FC (ca. 15% overestimation).

**Figure 2:**
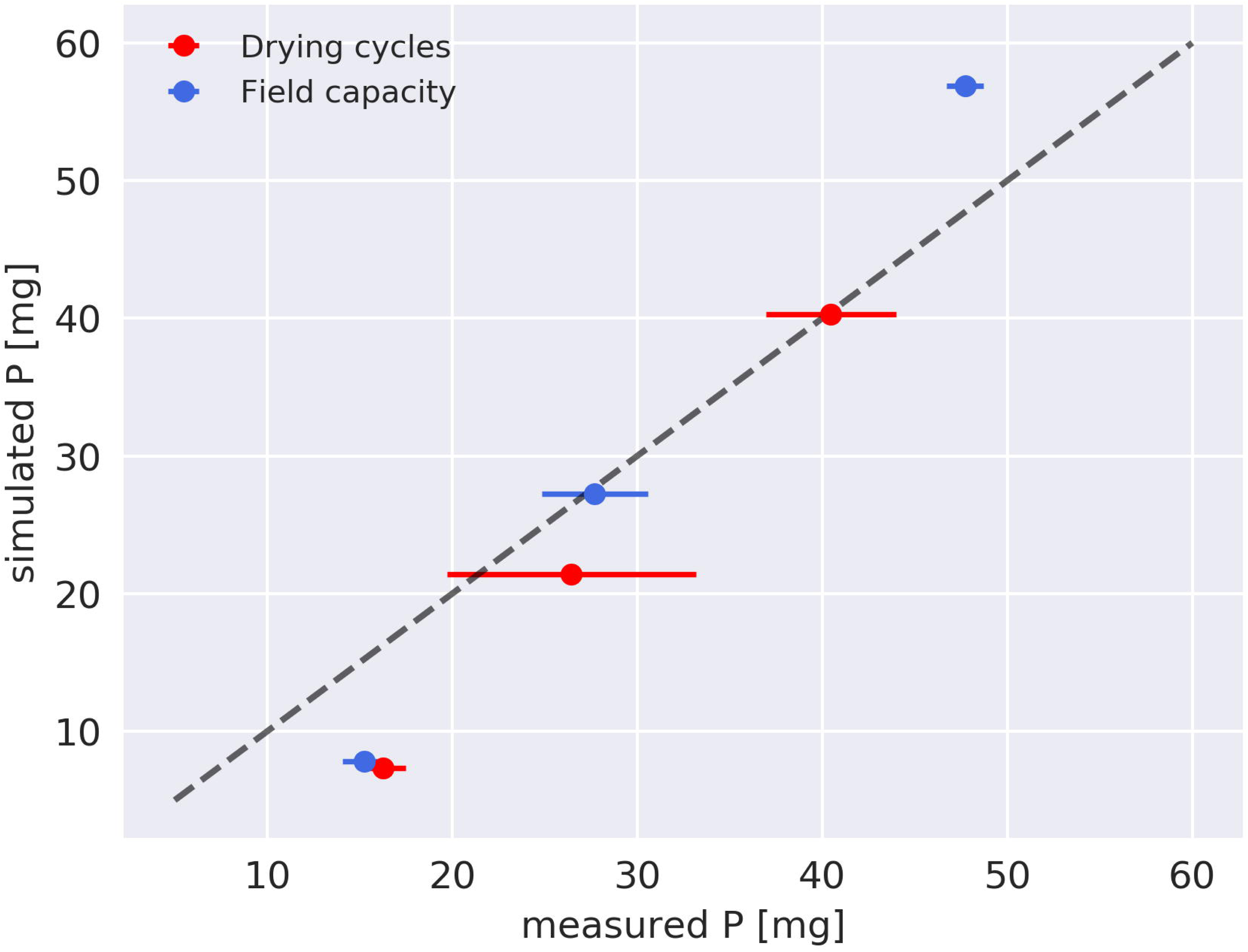
The simulated P uptake per plant (mg/plant or mg/pot, 1 plant per pot) versus the measured P uptake per plant in the lab experiment (including the standard error from the mean). The latter value of total P uptake was calculated for the shoot and the root after measuring shoot P concentration, and assuming an equal P concentration in the root. Rice root systems were grown and simulated on a P deficient soil with three P treatments in the topsoil (No P amendment (NoP), a suboptimal rate (SubP), and a non-limiting rate (PlusP)) and two water regimes (Field Capacity (FC) and Drying Periods (DP)).

### Determining the location of the water and P uptake through the multiscale model

The cumulative water uptake by rice in the pot experiment increased with increasing P rate and it decreased under DP. Measured water uptake by the root systems was used as a boundary condition in the model. However, simulations of the water uptake reveal that most water was generally taken up in the topsoil. Under FC ca. 70% of the total water uptake was acquired from the topsoil at 52 DAS. Interestingly, the relative contribution of the subsoil to water uptake increased under DP as this share increased to ca. 37% (compared to ca. 30% under FC). (Figure 3)

**Figure 3:**
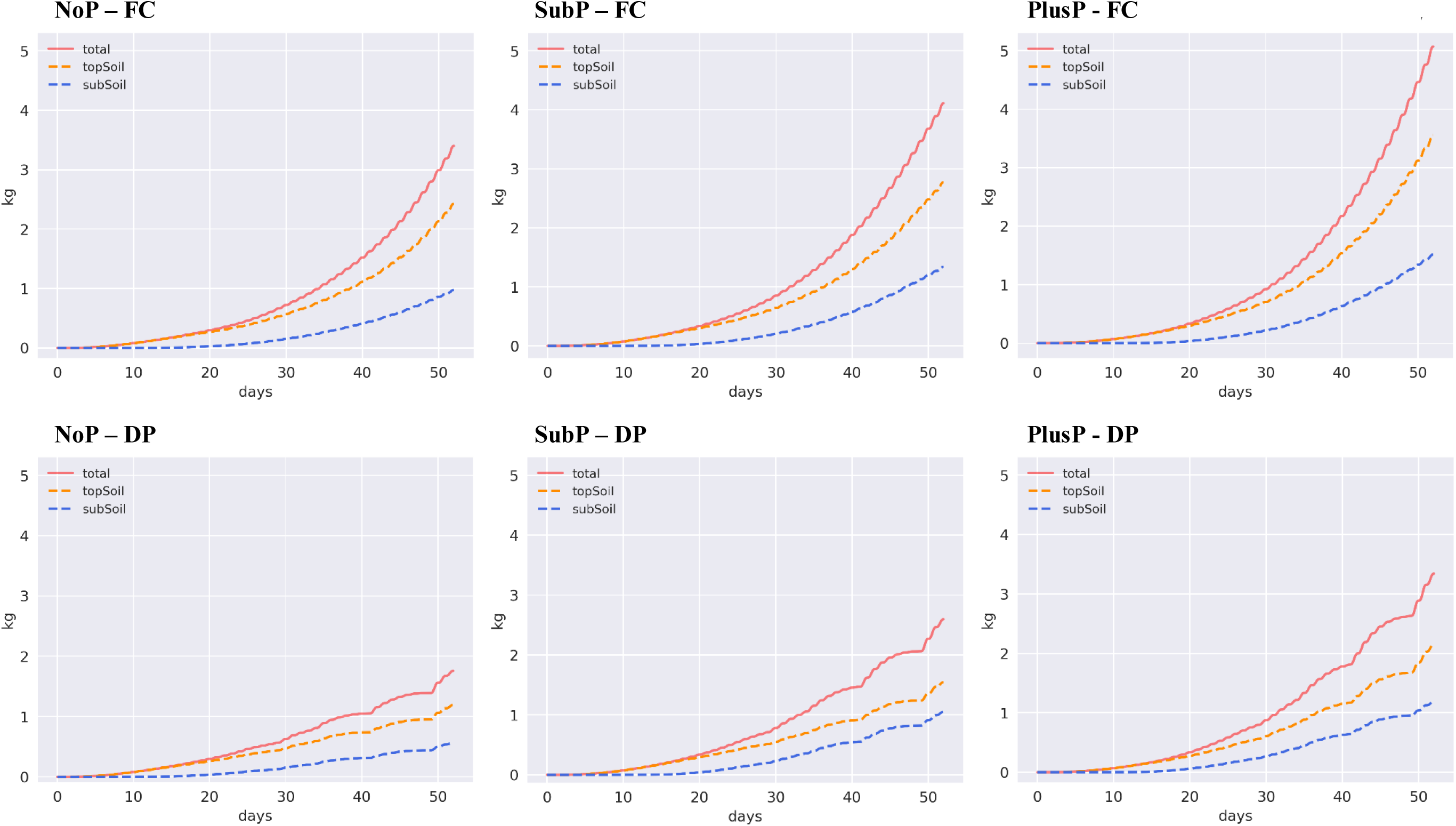
The simulated cumulative water uptake per plant (in kg) of rice roots during a growing period of 52 days. Rice roots were grown under contrasting P rates (P deficiency (NoP), a suboptimal P amendment in the topsoil (SubP), and a non-limiting P rate in the topsoil (PlusP)) and contrasting water regimes (field capacity (FC) versus drying periods (DP)). These simulations enable the differentiation among total water uptake, water uptake from the topsoil (0-20 cm depth), and water uptake from the subsoil (>20 cm depth).

Similar to water uptake, simulations reveal that P uptake was mainly located in the topsoil, also when the P availability in top- and subsoil were equally low (NoP; Figure 4). With increasing P rate, the percentage of P acquired from the topsoil increased (71% under NoP, 75% under SubP, and 88% under PlusP at 52 DAS, FC). In the scenarios without P application (NoP), the P uptake rate in topsoil was equal to the uptake rate in the subsoil (illustrated by the parallel lines on Figure 4). Interestingly, P acquisition from the subsoil slightly increased under DP (an increase in the contribution of ca. 10%), while the acquisition in the topsoil decreased (Figure 4). Figure S7 (supplementary information) illustrates the relation between the P uptake and the total water uptake. Interestingly, both experimental data and simulation results reveal that the lower water uptake under drying cycles has proportionally only a minor effect on the total P uptake.

**Figure 4:**
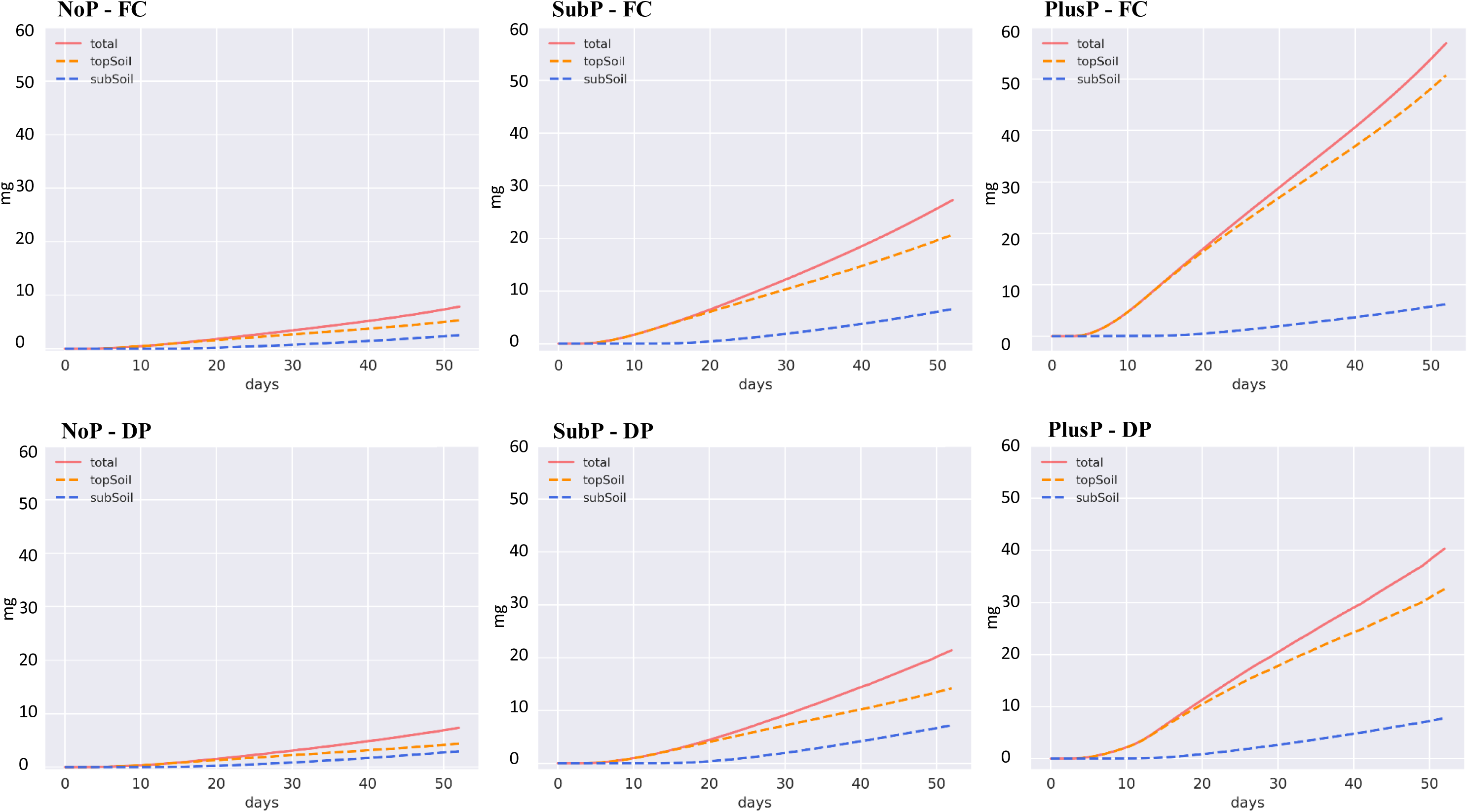
The simulated cumulative P uptake per plant of upland rice roots during a growing period of 52 days. Rice roots were grown under contrasting P rates (P deficiency (NoP), a suboptimal P amendment in the topsoil (SubP), and a non-limiting P rate in the topsoil (PlusP)) and contrasting water regimes (field capacity (FC) versus drying periods (DP)). These simulations enable the differentiation among total P uptake, P uptake from the topsoil (0-20 cm depth), and P uptake from the subsoil (>20 cm depth).

### The role of different root types and root characteristics for P and water uptake

Figure 5 displays the simulated contributions of each root type to water uptake. Under FC, the largest share in water uptake was attributed to S-type roots (47-59%). This trend altered under DP where the contribution of the L-type roots to water uptake (50-60%) exceeded the contribution of S-types (31-37%). Nodal roots consistently displayed only minor contribution to water uptake (Figure 5) and they showed a minor cumulative water uptake per unit root mass (Figure 7).

**Figure 5:**
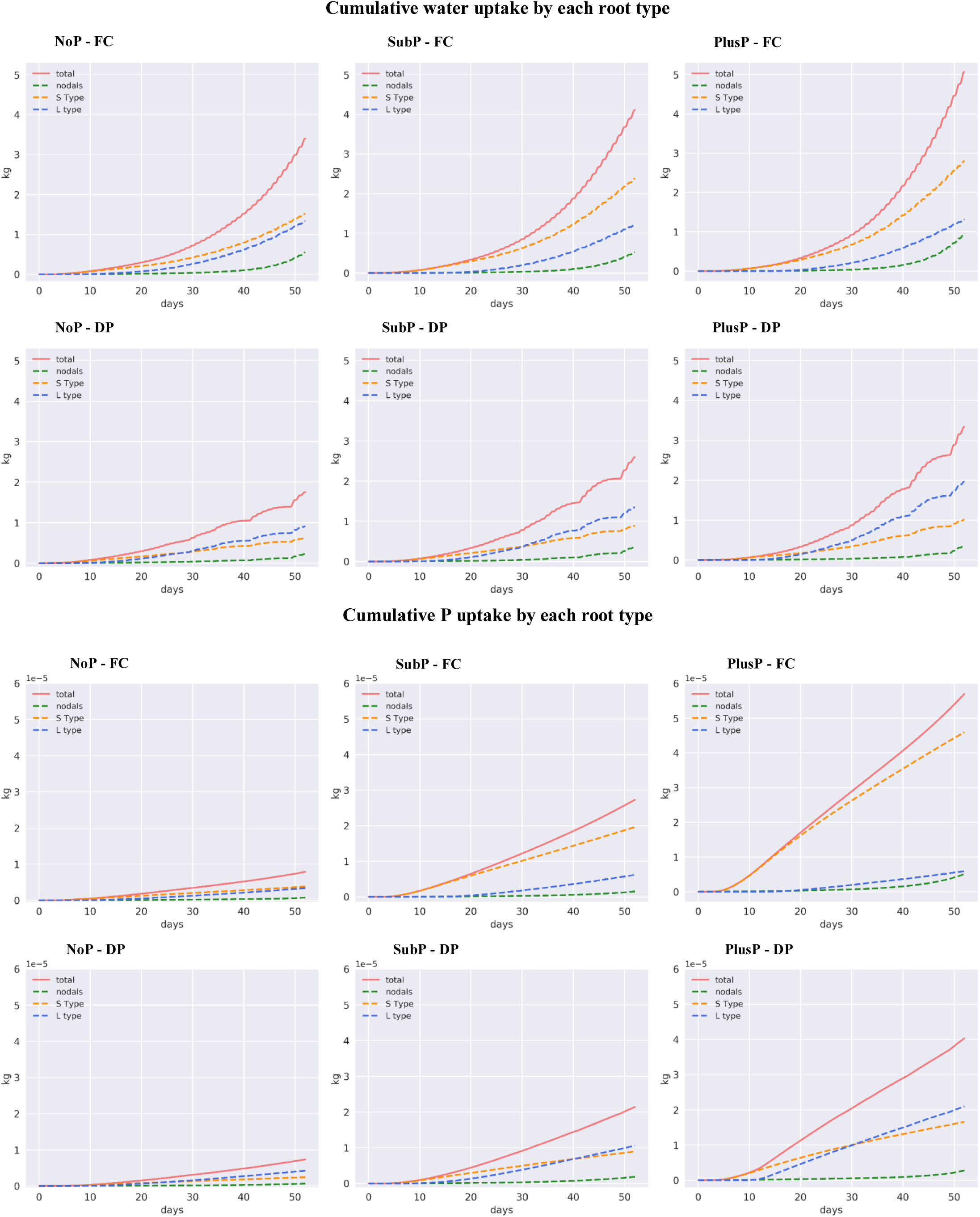
The cumulative water uptake per plant (top) and cumulative P uptake per plant (bottom) by the different root types of rice during a growing period of 52 days. Rice roots were grown under contrasting P rates (P deficiency (NoP), a suboptimal P amendment in the topsoil (SubP), and a non-limiting P rate in the topsoil (PlusP)) and contrasting water regimes (field capacity (FC) versus drying periods (DP)). These simulations enable the differentiation among the water uptake by the nodal roots, the S-type lateral roots, and the L-type lateral roots.

**Figure 6:**
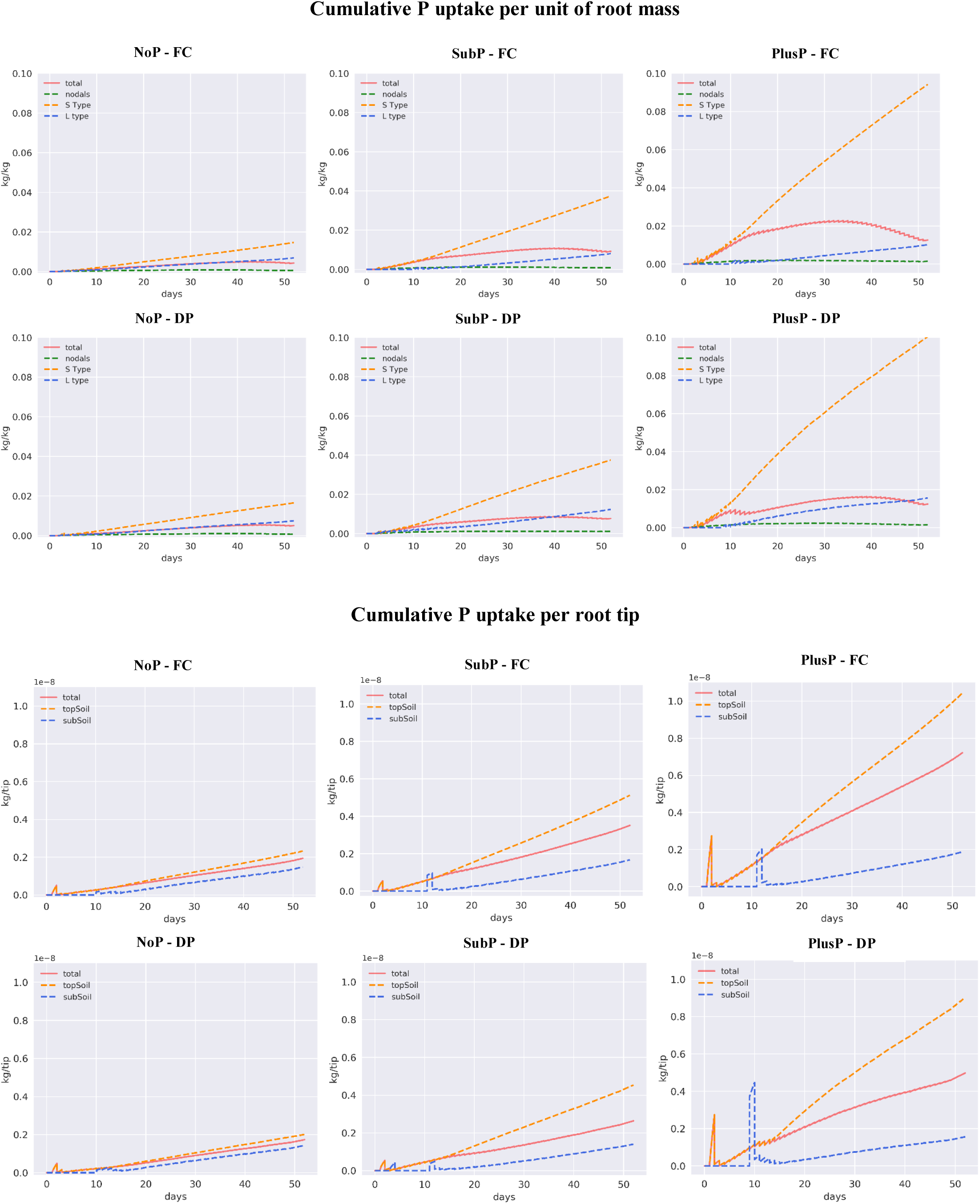
The simulated cumulative P uptake per unit of root mass for each root type (top) and the cumulative P uptake per root tip in each soil layer (bottom). When root tips start entering the subsoil (ca. 10 days), segment coordinates may be assigned to the subsoil, while a large part of the segment is still located in the topsoil. Therefore the P uptake is larger due to a larger availability, and hence peaks can be observed.

**Figure 7:**
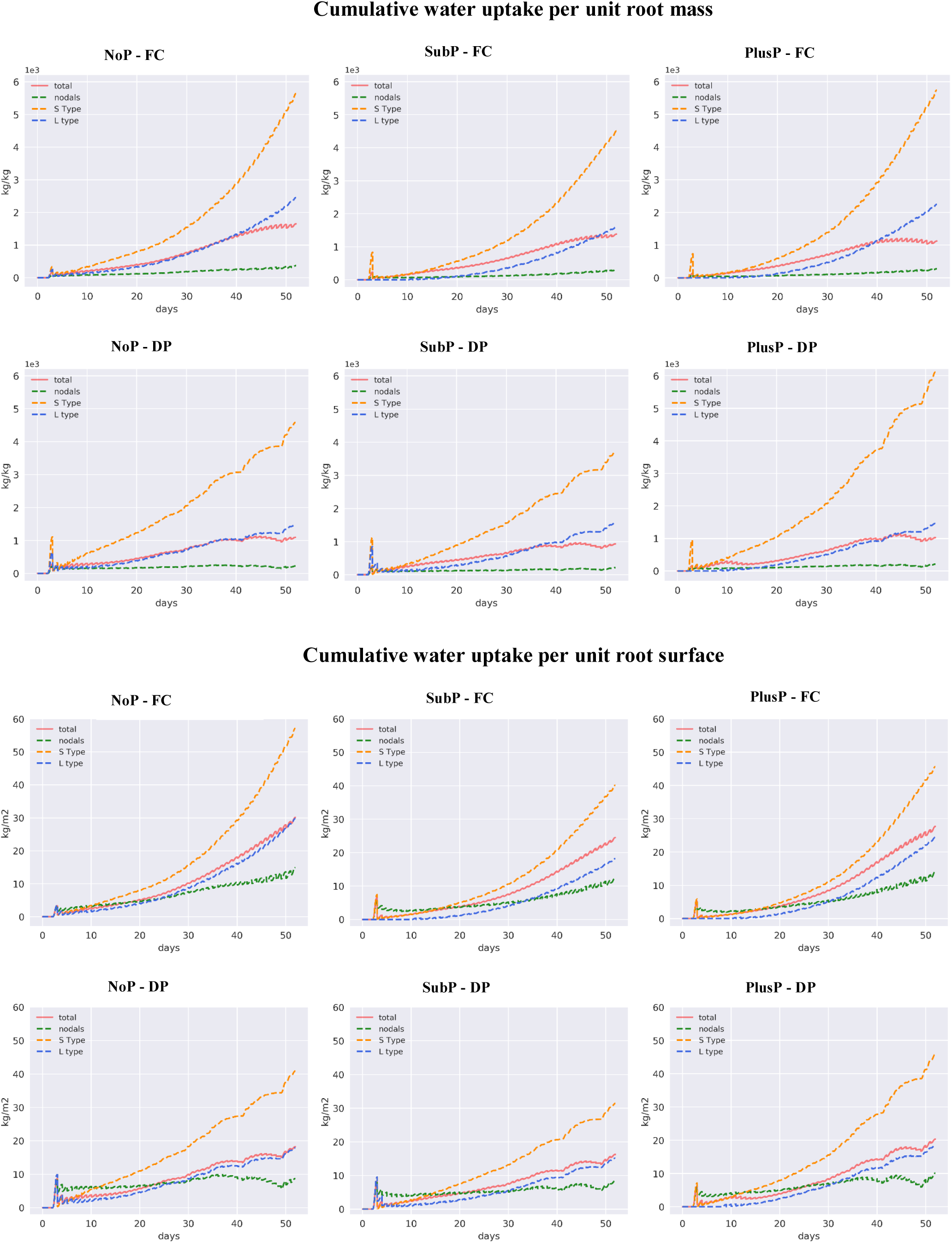
The simulated water uptake per unit of root mass for each root type (top) and water uptake per unit surface in each layer (bottom).

Similar to water uptake, simulations show that nodal roots generally have only minor contribution to P acquisition while S-type laterals displayed the largest contribution to P uptake under FC (Figure 5). Under DP, the contribution of the L-type roots to P uptake increased during the growth and the development of the root system and its contribution to P acquisition finally overruled the contribution of the S-type roots during development. With increasing P rates, the fractional contribution of the S-type roots to P uptake increased under both FC (from ca. 44% under NoP to 79% under PlusP) and DP (from ca. 31% to 44%), while the reverse was true for L-type roots. The cumulative P uptake per unit root mass equally increased for S- and L-type roots under NoP, while this ‘efficiency’ strongly increased for S-type roots under SubP and PlusP (Figure 6).

### Normalizing P and water uptake to root mass, root surface, root length, and root tips and the importance of root elongation

Under P limitations (NoP), a larger root mass was obtained under FC compared to the root mass under DP (Table 3). However, this larger root mass under FC did not lead to a large increase in P uptake (Figure 6), while this indeed resulted in a much larger increase of water uptake under FC compared to DP (Figure 7).

In contrast, under optimal P availability (PlusP) the larger root mass under FC compared to DP (Table 3) did lead to a larger increase in P uptake whereas the increase in water uptake per increase in root mass was smaller (Figure 6 & Figure 7).

At the end of the simulation period, the cumulative P uptake per unit of root mass (Figure 6) started to decrease in all scenarios. This trend is also observed for the cumulative P uptake per surface (not shown), however to a smaller extent. Both observations indicate a decline in P uptake rate per unit of root mass or surface. In contrast, for the cumulative P uptake per number of root tips one can observe a consistent linear increase towards the end of the simulation period, which indicate a constant P uptake rate per root tip (Figure 6), which is additionally supported by the findings displayed on Figure S6 (supplementary information). Figure S6 shows a general low P uptake rate by a single nodal root under NoP (Top) compared to SubP (Middle) and in the latter case it shows a sudden decrease in P uptake rate as soon as the root penetrates into the P deficient subsoil. Interestingly, the P uptake rate at the tip is generally large, but it dramatically decreases when the root does not continue to grow (Middle versus Bottom plot in Figure S6).

For water uptake the trends are different. The cumulative water uptake per root surface area still increases by the end of the simulation period, and the uptake rate remains relatively constant (with exceptions during the periods of water stress where the uptake per unit area decreases). This indicates a consistent increase in water uptake with increasing root surface. (Figure 7)

## Discussion

### Integrating the responses of root system architecture through 3D modelling

This work shows how 3D root architecture models such as CRootBox can be utilized to acquire insight in integrated root system responses. Multiple single root phenes (e.g. the number of nodal roots, nodal radius, lateral density, etc.) all contribute to the final performance of a root system but the utility of a given root phene for resource capture will often depend on the expression of other phenes. A large number of potential interactions, especially considering that synergism or antagonism among root phenes for soil resource acquisition will depend on environmental conditions, makes this a challenging research problem and therefore studies on an integrated root performance are generally scarce. It was previously argued that such functional-structural models are the only practical way to assess the large number of root phene interactions with other phenes or environmental variables through integrated systems and such an approach was therefore suggested as a useful tool to guide the breeding of ideotypes adapted to specific soil environments (Lynch 2011, 2019; Ahmadi *et al*. 2014; Rangarajan *et al*. 2018). Here we indeed demonstrate how such a functional-structural model can be used to integrate multiple co-occurring root phene responses and how it enables the evaluation of the total root system performance grown in several environments. This is assessed by calculating holistic root parameters that result from combining multiple single phenes (including both qualitative as quantitative root parameters) and that directly link to water or nutrient acquisition (e.g. total surface, surface distribution, number of root tips, total root length).

Clearly, this integrated approach permits a comprehensive evaluation of co-occuring responses of soil-grown rice root systems in the two-factorial landscape of P and water availability. It enables the identification of important phenes that contribute to tolerance under contrasting scenarios, as was previously argued by Lynch and Brown (2012). These simulations reveal how combined responses in root mass, nodal root number, root radii, and branching density, contribute to a decreased total root surface under decreasing P rate or water level, and it shows how the contribution of certain root types in upland rice root systems becomes more or less important in contrasting scenarios. In this regard, it is interesting that the total number of root tips and total root surface in the subsoil generally increased under drying periods compared to field capacity, and this seems to occur independently from P availability. Responses of single root phenes (such as deep root ratio, root angle, and branching) to drought events were indeed previously observed for upland rice (Kato *et al*. 2006; Gowda *et al*. 2011; Henry 2013; Menge *et al*. 2016), but to our knowledge this is the first study demonstrating the holistic, integrated responses of the total root system architecture.

A more efficient root system is generally attributed to roots displaying a high functioning with a relatively low cost (Lynch and Ho 2005). Generally, ‘root system efficiency’ was regarded as roots with a large surface to volume ratio, as soil exploration can hence occur at a lower carbon cost. With this perspective, this modeling study illustrates how multiple root phene responses (i.e. number of nodal roots, nodal diameter, S-type radius, L-type radius, S-type distance, and secondary distance) contribute to the functionality of a root system under suboptimal P availability (demonstrated by the high ‘surface/volume ratio’), and how this ‘root system efficiency’ generally improves under drying periods compared to field capacity. These observations correspond to the general theory of ‘rhizoeconomics’ (Lynch and Ho 2005; Lambers *et al*. 2006), and we have now proven that upland rice indeed reduces the cost of soil exploration by optimizing its root system performance (i.e. increased surface/volume, but also an increased root tip/volume ratio) under suboptimal P availability and drying periods by simultaneously altering multiple single root phenes. Interestingly, our study highlights the importance of growing root tips for P uptake in deficient soils rather than root surface only, and hence we argue that root functioning and root efficiency should be assessed by the parameters that actually drive the particular resource uptake in a particular condition. Obviously, ‘root efficiency’ would then depend on the specific characteristics of the stress conditions. In terms of P uptake this study thus highlights that root system efficiency should better be assessed by the ratio of emerging root tips over the total soil volume, while root system efficiency in terms of water uptake could indeed best be assessed by the surface-volume ratio, as discussed above. This complexity of different drivers in ‘resource uptake’ highlights the need of simulations to assess ‘root system efficiency’ in relation to multiple stresses. As such, we demonstrate that an increased investment in the root system in the subsoil towards the end of the simulation period led to similar or even lower P uptake rates from the subsoil versus the topsoil, while a higher uptake of water was observed from the deeper soil layer. The latter is attributed to the contrasting parameters affecting P or water uptake but it also follows a contrasting distribution of both resources in the soil.

### Insights from the 3D continuum multiscale model for phosphorus uptake by upland rice under contrasting water regimes

We have demonstrated how the coupled 3D continuum multiscale model of Mai *et al*. (2018) can be applied to simulate nutrient uptake from any crop grown in specific environments. Given the fact that there was no calibration of P dynamics, this model nicely simulated the P uptake for both water regimes (validation shown in Figure 2), and it presents the dynamics during root development, but also under drying periods. It is thus possible to simulate and unravel effects of drying periods on the P uptake of a root system grown in a highly P-fixing soil, and we demonstrate how these effects interact with the nutrient availability along depth.

Without P amendments (NoP), the P uptake was not influenced by the drying cycles, whereas water uptake was indeed affected by drying cycles. Under sub-optimal and optimal P conditions, drought stress also triggered a lower P uptake. It is interesting that the simulations strongly underestimated the P uptake only in the scenarios without P application, which can be explained by four factors. Firstly, under P deficiency the interaction of roots with the soil microbiome (e.g. mycorrhyzae) becomes highly important for soil exploration and this is not included (Maiti *et al*. 2011, 2017). To tackle this issue in modeling, a Matlab version of RootBox exists which considers primary and secondary infection of a growing root system with arbuscular mycorrhizal fungi (Schnepf *et al*. 2016), thus as an outlook, this can be coupled to the approach presented here. Secondly, root exudates may mobilize immobile P under P deficiency in order to enhance acquisition (Kirk *et al*. 1999; Schnepf *et al*. 2012; Tawaraya *et al*. 2013; Bhattacharyya *et al*. 2013). Thirdly, root hairs might play a key role on P uptake under limited availability (Leitner *et al*. 2010; Nestler and Wissuwa 2016), and these are not included in the root model. Hence, it should be considered that these simulations only assess physical aspects and therefore they might underestimate the actual P (or other nutrients) uptake when being deficient in soils. Lastly, the high root absorbing power for P (ratio of uptake flux to free ion concentration in solution; (Nye and Tinker 1969) suggests that diffusion-limited plant uptake is likely for P, even in nutrient solutions. As a consequence, the Michaelis constant, Km, derived from nutrient solution or sand cultures (as used here) are overestimated when based on the bulk solution ion concentration (Winne 1973). The Km values determined under diffusion limitations are therefore apparent values reflecting the physical process (diffusion limitation), and are not characteristic for the biological transport process (transporter affinity towards the transported ion). Santner *et al*. (2012) argued that the true Km values for plant root P transporters would be much lower than the values usually reported in literature. For models where Michaelis–Menten kinetics is used, they recommended to use a low value for Km (e.g. the one estimated in buffered solutions: Km ∼ 0.5 µmol l^−1^) rather than a value determined in unbuffered solutions. As the Km value used in this model was determined on gravel and nutrient solution (Teo et al. 1992b), diffusion limitation might have occurred and hence we re-simulated our model for two scenarios with a smaller Km value as suggested by Santner *et al*. (2012) (i.e. 0.5 µmol l^−1^, which is 7 times smaller than the Km value used in this study), and also using a smaller Vmax (i.e. 1.84E-10 kg m^−2^ s^−1^; which was determined by using the average uptake rate initially simulated in the PlusP-FC scenario from which the interface concentration was derived. The new Vmax was derived so that for this interface concentration and new Km value, the Michaelis-Menten uptake reproduced the same uptake rate. When using this small Km and smaller Vmax, the simulated P uptake under the NoP scenario strongly improved (Figure S12) while the simulations under SubP and PlusP barely changed. This highlights the need of using buffered Km values when simulating root P uptake under deficient conditions and it might indeed suggest that the general Km values for P uptake currently available in literature are too large (Santner *et al*. 2012).

### Effects of integrated root system responses on local P and water acquisition, and drivers in uptake

This study demonstrates how this coupled structural-functional model allows interpretations on the importance of soil resource uptake in certain soil layers, and it allows quantification along the depth. Indeed it was previously suggested for upland rice that water is preferentially extracted from the top layer and that more water is taken up from deeper layers only after drying of the topsoil (Kondo *et al*. 2000; Price *et al*. 2002), however, to our knowledge this was never quantified before. Deep rooting enhances drought tolerance by enhancing deep water acquisition (Fukai and Cooper 1995; Uga *et al*. 2013; Lynch and Wojciechowski 2015), and deep water acquisition was generally regarded as a drought adaptation in crops (Comas *et al*. 2013). Interestingly, this work now reveals that ‘deep water acquisition’ in upland rice only comprised a relatively small fraction of the total cumulated water uptake, even under droughts. It should however be noted that water stress only occurred during the second half of the experiment and impact of deep roots could thus only be revealed at the later stage. Interestingly, at the end of these simulations, the water uptake rate from the subsoil indeed became similar to the uptake rate in the topsoil. Hence we argue that, although the total fraction of water uptake over the total growing period from the subsoil is small, deep roots indeed contribute to plant survival under water stress. Therefore, future research should thus utilize this 3D continuum multiscale model to identify root ideotypes or root parameters that can further enhance deep water acquisition as also discussed by Asch *et al*. (2005).

Similar for P uptake, we quantified the importance of P acquisition from the topsoil. Topsoil foraging of P generally follows from the P accumulation in upper layers (Fei *et al*. 2011; Lal and Stewart 2016), but this study shows that this trend also holds when P availability is equally distributed in a soil column, even when a larger root surface is found in the subsoil. The latter trend can only be explained by the longer residence time of roots in top layers (grown from root base to bottom), and topsoil foraging of P thus not only follows from a higher P availability in upper layers or the final root surface distribution. Additionally, this work highlights that P uptake from deeper layers by upland rice should not be ignored, especially when grown in soils subjected to droughts ‘deep P acquisition’ increases. Interestingly, by increasing water availability in a P deficient soil, transpiration is increased, but hardly P uptake, and this would lead to a lower WUE. It is shown that water uptake per unit root mass increases with increasing water application whereas the P uptake per unit root mass decreases with increasing water application/uptake in a P deficient soil.

Our simulations indicate a general depletion of P along the root axis which results in a reduced P uptake rate from the tip towards the basal end of the roots (Figure 6, S5). With P applied in the topsoil (SubP and PlusP), a decrease in P uptake per root mass or surface would also be a consequence of more new roots that develop in the deeper P deficient soil layer. These simulations highlight the importance of root tips and root elongation for P uptake and we argue that increasing the length of a single root will not tremendously increase the P uptake but that the emergence of new roots will lead to a stronger increase in P uptake. Several studies have found that the regions close to the root tip can be biochemically more active, having a higher density of phosphate transporters (Smith 2002; Péret *et al*. 2011; Kanno *et al*. 2016), or following after the development of aerenchyma starting from a few centimeters above the root tip in rice. We show that the higher importance of the root tips to overall P uptake can also be explained by physical principles alone, i.e., due to the high depletion of P near the root surface, root tips that grow into still undepleted soil have higher contribution even when the same amount of transporters/same Michaelis Menten parameters are prescribed along the root axis. In our model, we assumed that the ‘uptake capacity’ of a root did not change along the root, but we show that this uptake capacity becomes ‘latent’ behind the root tip because of P transport limitations in the soil. Thus, it would not make sense for a plant to have many transporters near the base of the root.

In the rhizosphere model, uptake of nutrients will create a depletion zone around each root segment in which the gradient of nutrient concentration is largest at the root surface. We defined the depletion radius as the distance from the root center to the position where the nutrient concentration gradient decreases to 1% of that at the root surface. Comparing with the halfmean distance values, the depletion radii (Figure S8) are generally two times smaller than the calculated half mean distance in the NoP scenario (Table 3), while the halfmean distance and the depletion radii are comparable in the PlusP scenario. This indicates that competition between roots for P uptake occurs more likely in the PlusP scenarios due to the large depletion radii and large root density in the soil. Furthermore under NoP and SubP, P uptake by newly developed root tips will not be influenced by the P uptake of the other root segments or by depletion of the P stock in the soil by previous uptake. These findings suggest that one should develop rice varieties developing root systems with many branches and many new emerging tips in order to enhance low P tolerance in rice, rather than creating varieties with a large root surface area only (Rose *et al*. 2013).

Interestingly, such root systems showing a high branching density with many root tips would also be beneficial for root penetration and this would also improve the plant’s tolerance to drying periods (Bengough *et al*. 2011).

Our study enables to differentiate the contribution of particular root types for P and water uptake, and it explains how integrated root responses influence soil resource acquisition. Indeed, we demonstrate that nodal roots have only minor contribution to water or P uptake and nodal roots thus rather function as the ‘skeleton’ determining the spatial distribution in soil (Fageria 2013), but a reduced nodal thickness might indeed contribute to a more efficient biomass utilization. S-type roots are found to have a large contribution to P uptake, and De Bauw *et al*. (2019) previously related a higher density of such S-types with an increased P uptake capacity under deficient conditions. Interestingly, the upland rice variety simulated in this study (i.e. NERICA4) was previously found to have a very sparse S-type density compared to other varieties (e.g. NERICA-L-19, Mudgo, & DJ123) and the role of S-type roots on P uptake would thus be even more important for these other varieties. In a next phase, this model can be used to investigate whether the contrasting perfiormance of other varieties more tolerant to drought or low P can be explained by their root system characteristics. This study additionally highlights that the P uptake by L-type roots should definitely not be ignored in upland conditions as their contribution in P uptake even overrules the uptake of S-type roots under droughts. We mathematically demonstrate that the previously observed increased P uptake efficiency in response to reduced water availability (De Bauw *et al*. 2019) mainly attributes to an increased P uptake by L-type roots following a higher secondary branching density. The latter drought adaptation of upland rice indeed enhances a more efficient water (and P) uptake through an improved functioning of L-type roots. Previous studies highlighted the importance of reduced lateral branching to drought tolerance (Gao and Lynch 2016), but our simulations demonstrate that major part of the water under field capacity is acquired by S-type roots and that indeed the water uptake by L-type roots also increases under droughts. In search for root traits that benefit growth under both P deficiency and drought stress, no characteristics could previously be determined, however, this study now indicates that the key to a synergistic tolerance to droughts and low P can be found in the secondary branching of L-type roots.

## Conclusions

This work shows how in-silico experiments after model-data integration enhance our mechanistic insights in soil-root processes. It demonstrates how multiple co-occuring single root phene responses to environmental stimuli such as water and P availability contribute to the development of a more efficient root system under drying cycles or sub-optimal P availabililty.

General drivers in soil P and water uptake are unraveled and we describe how these alter under contrasting scenarios. For P uptake under deficient conditions, we demonstrate the importance of growing root tips. Additionally, this study allowed the quantification of the contribution of contrasting root types to both P and water uptake and the most relevant root characteristics that enhance low P and/or drought tolerance were identified. The model would need some further improvements to adjust for the low simulated P uptake under deficient conditions. To this extent, modelers should consider the use of a Km that is derived from solutions in which the low P concentrations are buffered and models may be refined by including the effects of root exudates, root hairs, and infections of mycorrhizae.

## Funding information

The pot trial and lab analyses were partly conducted at and financed by the Africa Rice Center in Tanzania and the KU Leuven in Belgium, and the work was additionally supported by the Belgian VLIR-UOS through a scholarship (VLADOC grant awarded to P. De Bauw). T.H. Mai was funded by the German Federal Ministry of Education and Research (BMBF) in the framework of the funding initiative “Soil as a Sustainable Resource for the Bioeconomy BonaRes”, project “BonaRes (Module A): Sustainable Subsoil Management - Soil3; subproject 3” (grant 031B0026C).

## Acknowledgements

We thank Dr. Elke Vandamme en Dr. Kalimuthu Senthilkumar for their hospitality and assistance at the Africa Rice Center in Tanzania and we thank Allen Lupembe and Nassoro Hemedi for the maintenance of the pot trial.

## Captures Supplementary Files

Figure S1: The observed root mass distribution along depth (A: 0-15 cm; B: 15-30 cm; C: >30 cm) in the pot experiment (colored bars including the standard error) and the corresponding distribution simulated in CRootBox (black triangle) in all scenarios.

Figure S2. Boundary conditions for six in-silico experiments of the virtual soil-root system. The daily evaporation and irrigation rate (left) and the prescribed transpiration rates at root collar (right).

Figure S3: The interbranch distance scaling factor along the soil depth. This figure shows the scaling factor of the interbranch distance of the laterals on nodal roots, following the particular root mass distribution in each scenario.

Figure S4: The probability of the S-type roots along the soil depth. This figure shows a high probability of the S-type laterals starting to emerge from the nodal roots in the top soil until the transition depth. Below the transition depth and in the subsoil layer, there are no S-type laterals (zero probability), but only L-type laterals emerging from the nodal roots. The lines for SubP-FC and PlusP-FC overlap on the figure.

Figure S5: The simulated root systems from CRootBox showing the different root types (nodal roots = red; S-type roots = yellow; L-type roots = green) and the corresponding emergence.

Figure S6: The simulated P uptake rate along a nodal root (kg s-1 m-1) versus the distance from the root origin for (i) a growing root under NoP conditions (Left), (ii) a growing root under SubP conditions (Middle); (iii) a non-growing root under SubP conditions (Right).

Figure S7: The relation between water uptake and the measured (left) and simulated (right) P uptake for each scenario.

Figure S8: Depletion radii in the rhizospheres (Top) and the P uptake rates of root segments in the rhizospheres (Bottom) for each scenario during the simulation period of 52 days. The center line represents the mean value, while different color shades represent the 25-75 percentiles (interquartile range) and the 5-95 percentiles of the root segment uptake rates.

Figure S9: The simulated cumulative P uptake per unit of root length for each root type.

Figure S10: The dynamics of the water content (cm3 cm-3) in the soil during the simulation period.

Figure S11: The effective P diffusion coefficient (m²s-1) at the root surface (extracted from rhizosphere models). Under DP, the diffusion coefficient clearly drops during drying.

Figure S12: The re-simulated P uptake per plant versus the measured P uptake per plant in the lab experiment (including the standard error from the mean) by using a small Km (1.55E-05 kg m^−3^) and a small Vmax (1.84E-10 kg m^−2^ s^−1^). The measured value of total P uptake was calculated for the shoot and the root after measuring shoot P concentration, and assuming an equal P concentration in the root. Rice root systems were grown and simulated on a P deficient soil with three P treatments in the topsoil (No P amendment (NoP), a suboptimal rate (SubP), and a non-limiting rate (PlusP)) and two water regimes (Field Capacity (FC) and Drying Periods (DP)).

Supplementary text: Detailed information on the model descriptors and the mathematical equations

Supplementary Video 1: The simulated root growth, P uptake rates (color scales on the root), and soil P sink (color scales in the rhizosphere), during the growing period of 52 days for the scenario under Suboptimal P and Field Capacity.

